# Speech sound categorization: The contribution of non-auditory and auditory cortical regions

**DOI:** 10.1101/2021.10.08.463391

**Authors:** Basil Preisig, Lars Riecke, Alexis Hervais-Adelman

**Author notes:** Address for correspondence: Basil C. Preisig, PhD.

## Abstract

Which processes in the human brain lead to the categorical perception of speech sounds? Investigation of this question is hampered by the fact that categorical speech perception is normally confounded by acoustic differences in the stimulus. By using ambiguous sounds, however, it is possible to dissociate acoustic from perceptual stimulus representations. Twenty-seven normally hearing individuals took part in an fMRI study in which they were presented with an ambiguous syllable (intermediate between /da/ and /ga/) in one ear and with disambiguating acoustic feature (third formant, F3) in the other ear. Multi-voxel pattern searchlight analysis was used to identify brain areas that consistently differentiated between response patterns associated with different syllable reports. By comparing responses to different stimuli with identical syllable reports and identical stimuli with different syllable reports, we disambiguated whether these regions primarily differentiated the acoustics of the stimuli or the syllable report. We found that BOLD activity patterns in left perisylvian regions (STG, SMG), left inferior frontal regions (vMC, IFG, AI), left supplementary motor cortex (SMA/pre-SMA), and right motor and somatosensory regions (M1/S1) represent listeners’ syllable report irrespective of stimulus acoustics. Most of these regions are outside of what is traditionally regarded as auditory or phonological processing areas. Our results indicate that the process of speech sound categorization implicates decision-making mechanisms and auditory-motor transformations.

**Highlights:**

- Ambiguous dichotic syllables elicit distinct percepts of identical stimuli
- Multivariate searchlight analysis reveals syllabic-category sensitive brain areas
- Categorical responses arise in non-auditory cortical areas including motor areas
- SMA is a possible locus for transforming sensory signals into perceptual decisions

## 1 Introduction

The mapping of sensory information onto common categories is a fundamental feature of human cognition. For example, we can identify a familiar person on different photographs even if taken from different angles. Likewise, we can map speech sounds onto common words even if uttered from different speakers. Categorical perception in speech was described first by Liberman and colleagues (1957) who showed that synthetic syllables along the continuum between prototypes (e.g., /ba/ vs /da/) were perceived categorically despite their incremental acoustic variations. However, it remains unclear how the brain maps the large variety of sensory signals to a limited number of invariant categories.

Speech sound categorization can be conceptualized as the graded and continuous process from sensory perception to action. Consistent with this notion, previous studies have identified multiple auditory and non-auditory brain regions, where speech sound categories can be segregated (Myers et al., 2009). Several studies identified categorical phoneme representations in the auditory association cortex including the superior temporal gyrus (STG) and the superior temporal (STS) using fMRI (Formisano et al., 2008; Kilian-Hütten et al., 2011; Arsenault and Buchsbaum, 2015; Levy and Wilson, 2020) and intracranial recordings (Chang et al., 2010; Mesgarani et al., 2014; Yi et al., 2019). Other studies identified phoneme representations in the motor cortex using fMRI (Pulvermüller et al., 2006; Lee et al., 2012; Chevillet et al., 2013; Du et al., 2014; Evans and Davis, 2015) and intracranial recordings (Cheung et al., 2016). Evidence that is consistent with the role of motor cortical areas in speech sound categorization is provided by non-invasive brain stimulation studies which perturbed phoneme perception when applied directly over the motor cortex (D’Ausilio et al., 2009; Möttönen and Watkins, 2009; Smalle et al., 2015). Further studies found neural response patterns consistent with categorical encoding in frontal areas like the inferior frontal gyrus possibly involved in executive and decision-making processes (Hasson et al., 2007; Myers et al., 2009; Lee et al., 2012). Lastly, categorical responses to speech sounds have been detected in regions associated with phonological processing such as the left angular gyrus (Blumstein et al., 2005) and the left supramarginal gyrus (Caplan et al., 1995; Zevin and McCandliss, 2005; Raizada and Poldrack, 2007).

A major methodological challenge is to identify brain regions in which neural activity tracks the perceived speech rather than its sensory properties. For instance, if physically different stimuli are used as exemplars of different categories, the perceptual representation of the stimulus is confounded with its physical properties. Under such circumstances, it is thus difficult to disentangle whether observed neural responses represent the different stimuli, their different percepts, or both. One approach to overcoming this problem is to present two stimuli that are different at the level of sensory encoding, but result in the same speech percept (Formisano et al., 2008; Evans and Davis, 2015). In an fMRI study, Hasson and colleagues (2007) exploited the McGurk effect (Mcgurk & Macdonald, 1976) to present physically different stimuli (audiovisual /ta/ vs. auditory /ka/ + visual /pa/) that elicit the same phonemic percept (/ta/). Using this approach, they identified brain regions that show neural facilitation for stimuli with shared perceptual, but different sensory, encoding. This results were confirmed by Skipper and colleagues (2007) who showed that patterns in these frontal motor areas resulting from the illusory /ta/ percept are more similar to the activity patterns evoked by audiovisual /ta/ than they are to patterns evoked by audiovisual /pa/ or audiovisual /ka/. However, this approach does not easily allow differentiation between areas implicated in audiovisual integration from those involved in categorical speech perception. Another approach is to employ ambiguous stimuli that can elicit different percepts in order to identify brain regions that represent the perceptual report of the participant given the same acoustic stimulus (Kilian-Hütten et al., 2011; Lee et al., 2012). This approach, however, weights perceptual representations resulting from sensory uncertainty more strongly, which may not generalize well to situations with higher sensory evidence.

In contrast to previous research, the present study aimed to identify regions that consistently differentiate perceptual reports of both unambiguous and ambiguous stimuli. The advantage of this approach is that the identified activity patterns consistently differentiate the perceptual report in the presence and absence of an acoustic stimulus difference. For this purpose, we re-analyzed data from a previous fMRI study (Preisig et al., 2021) in which participants were presented with ambiguous (intermediate between /ga/ and /da/) or unambiguous stimuli (clear /ga/ vs. /da/) to right ear (RE) together with a disambiguating speech feature (high vs low third formant, F3) to the left ear (LE). In the ambiguous condition, the syllabic identity of the stimulus in the RE had been rendered indeterminate by shifting the F3 to the participants’ individual categorization boundary, i.e. the F3 level at which each participant reported perceiving the stimulus as /da/ or /ga/ in ∼50% of the trials during a pre-test (see Fig. 1A). The concurrent presentation of the disambiguating feature of this syllable (low F3 supporting a /ga/ interpretation or high F3 supporting a /da/ interpretation) to the LE can lead to an integrated syllable percept resulting in a classification consistent with the F3 stimulus presented to the LE. Please note that the F3 does not represent a meaningful speech sound on its own. In the unambiguous condition, the categorization of the syllable did not depend on the integration LE and RE input (see Fig 1B).

**Fig. 1.**
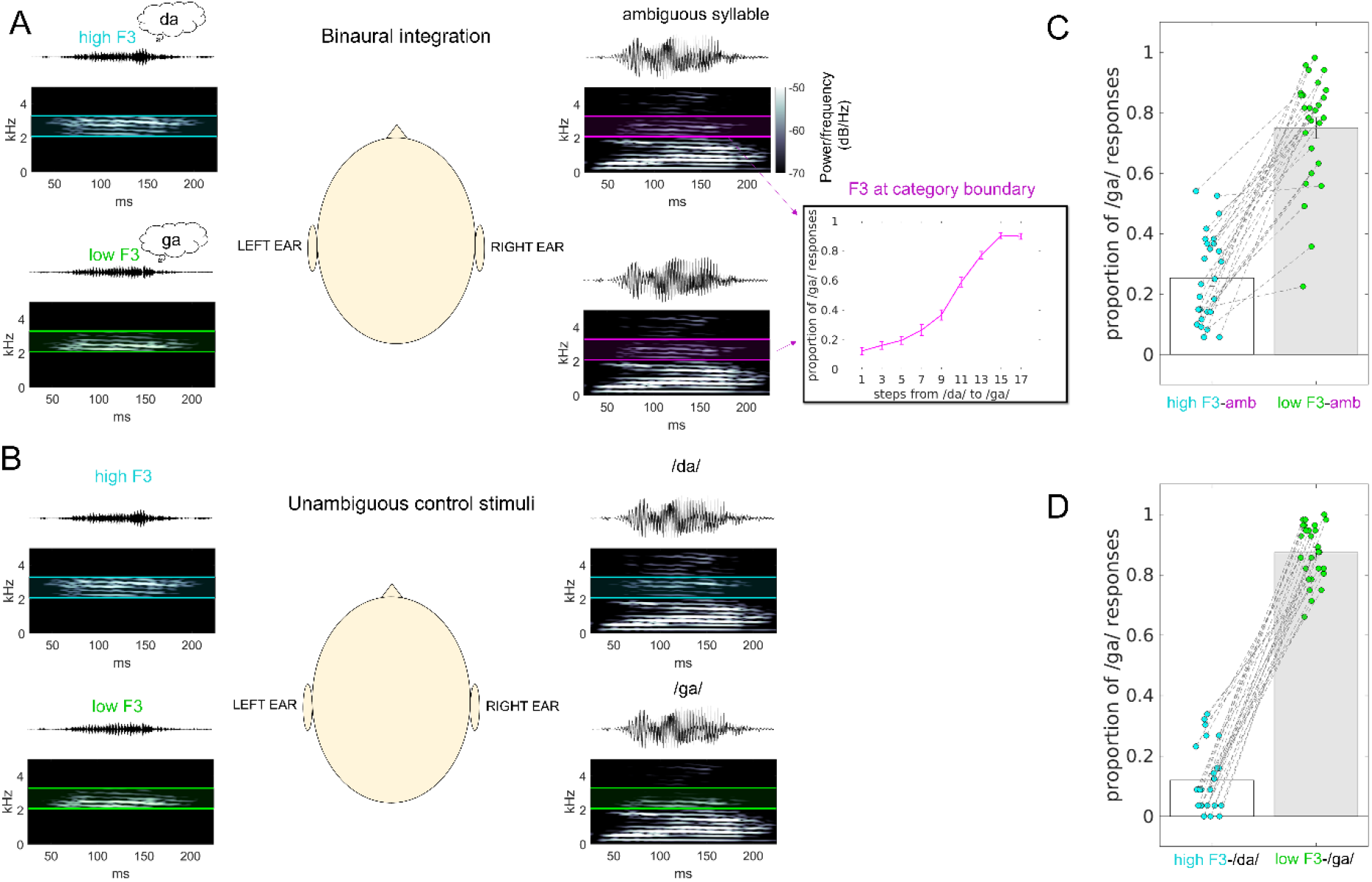
Auditory stimuli and behavioral responses. For each sound stimulus the pressure waveform is presented on top of the corresponding sound spectrogram (A) In the binaural integration condition, the perceived identity of the syllable depends on the integration of the right and left ear inputs. An ambiguous syllable (intermediate between /ga/ and /da/) was presented to the RE and a disambiguating speech feature (high vs low F3) was presented to the LE. Inset, the level of the F3 for the ambiguous syllable was determined in a pre-test as the intermediate step at which each participant reported perceiving the stimulus as /da/ or /ga/ in ∼50% of the trials. The figure shows the proportion of /ga/ responses across participants as a function of F3 step from /da/ to /ga/ (mean ± SEM). (B) In the unambiguous control stimuli, a clear /da/ or /ga/ stimulus was presented to the RE. The LE stimulus included a F3 cue with the same F3 frequency as the RE stimulus. (C) The proportion of /ga/ responses in the binaural integration condition as a function of the presented speech feature (high vs low F3). (D) The proportion of /ga/ responses in the unambiguous control condition as a function of the presented syllable (/ga/ vs /da/).

Multi-voxel pattern searchlight analysis (Kriegeskorte et al., 2006) was used to identify brain areas that consistently differentiated between response patterns associated with different syllable reports. Further, we tested whether the identified brain regions carry information about the perceived speech (/da/ vs /ga/ response) alone, or its acoustic properties (high vs low F3). Some previous fMRI studies have applied MVPA to investigate categorical speech perception (Formisano et al., 2008; Raizada et al., 2010; Kilian-Hütten et al., 2011; Lee et al., 2012). However, most of these studies have restricted their analysis to auditory cortical regions. Based on these previous results, we predicted that response patterns in auditory cortical regions consistently differentiate perceptual reports of both unambiguous and ambiguous stimuli. Using a searchlight procedure, we explored whether non-auditory cortical regions also show such differentiated patterns of response.

## 2 Material & Methods

### 2.1 Participants

Twenty-seven right-handed listeners with no history of hearing impairment (M=21.89 years, SD=3.14, 8 male) took part. The present analysis is based on a dataset collected as part of our previous research on the influence of transcranial brain stimulation on binaural integration (Preisig et al., 2021). All participants had normal or corrected-to-normal visual acuity. The participants reported no history of neurological, psychiatric, nor hearing disorders. All participants had normal hearing (hearing thresholds of less than 25 dB HL at 250, 500, 750, 1000, 1500, 3000, and 4000 Hz, tested on each ear using pure tone audiometry) and no threshold difference between the left ear (LE) and the right ear (RE) larger than 5dB for any of the tested frequencies. All participants gave written informed consent prior to the experiment. This study was approved by the local research ethics committee (CMO region Arnhem-Nijmegen) and was conducted in accordance with the principles of the latest Declaration of Helsinki.

### 2.2 Auditory Stimuli

The auditory stimuli were based on stimuli reported in previous work (Preisig and Sjerps, 2019; Preisig et al., 2020, 2021). A /da/ to /ga/ stimulus continuum was created in Praat software (Boersma and Weenink, 2019) by shifting the third formant (F3) in 17 equidistant steps from ∼2.9 to ∼2.5 kHz. The intermediate step at which each participant reported perceiving the stimulus as /da/ or /ga/ in ∼50% of the trials was selected as the ambiguous stimuli for RE presentation (see Fig. S1 for individual psychometric curves). To generate the F3 stimulus for LE presentation, the third formant (F3) was extracted from the endpoints of the continuum (from /da/ and /ga/) using a bandpass filter with frequencies between 2100 and 3300 Hz. For a schematic representation of the stimuli see Fig. 1A and B.

### 2.3 Experimental Design and Task

The dataset reported in this article comprised four task fMRI runs and one fMRI run with passive listening. The data from four additional task runs during which participants underwent non-invasive brain stimulation are reported elsewhere (Preisig et al., 2021).

Each task fMRI run comprised 128 trials, of which 88 trials included the presentation of an auditory stimulus (four trials were discarded because they occurred during a time period where non-invasive brain stimulation was turned on and off to evoke sensations associated with the initial onset and offset of stimulation). Auditory stimuli were presented using MR-compatible insert earphones (Etymotic ER-30) at a sound level of approximately 70 dB SPL. Each task fMRI run included 60 binaural integration trials, for which the F3 frequency of the RE stimulus was set at the individual category boundary (see Fig. 1A), and 24 unambiguous control trials, for which the F3 component of the RE stimulus supported 12 times a clear /da/ and 12 times a clear /ga/ interpretation (see Fig. 1B). For binaural integration trials, the LE stimulus was 30 times the high F3 and 30 times the low F3. For control trials, the LE stimulus included a F3 cue with the same F3 frequency as the RE stimulus. In control trials, perception of syllable category would be unaffected by binaural integration, because an unambiguous instance of the syllables /da/ or /ga/ was presented to the RE. It cannot be ruled out that participants used different listening strategies for different stimulus conditions, e.g., ignoring inputs to one or the other ear during the task. However, we consider it very unlikely that participants deliberately ignored input to one ear or the other because these stimuli are usually experienced as a fused acoustic image (the stimuli can be downloaded from our previous publication Preisig and Sjerps, 2019).

The less frequent unambiguous control trials were interspersed at random positions during the presentation of the binaural integration trials. Previously, we found a slight response time advantage for the LE (Preisig and Sjerps, 2019). Hence, the F3 was presented to the LE and the syllable stimulus (ambiguous or unambiguous) was presented to the RE.

Participants were asked to indicate by button press with the index finger of their left hand whether they heard the syllables /da/ or /ga/. Responses were recorded from the upper two buttons of an MR-compatible 4-buttons response-box (Current Design, HHSC-1×4-CL). The response pad was placed in the participants’ hand, and their index finger was placed on the relevant buttons.

During task fMRI runs, each trial was 3 s long (equal to the repetition time of the fMRI sequence) and started with the acquisition of a single fMRI volume (TA = 930 ms). The auditory stimulus was presented 1750 ms after the trial onset plus a pseudorandom interval (between 4 and 24 ms, in steps of 4 ms) (Preisig et al., 2021). The presentation of the auditory stimulus lasted 250ms. The participant’s response window corresponded to the interval from the auditory stimulus onset to 70 ms before the onset of the next trial.

The passive listening fMRI run consisted of 336 trials, of which 96 trials included auditory stimuli: 48 binaural integration trials and 48 unambiguous trials. The stimulus presentation order was randomized with the constraint that the transition rates between the four different stimuli (LE high F3 / RE ambiguous syllable; LE low F3 / RE ambiguous syllable; LE high F3 / RE clear /da/; LE low F3 / RE clear ga/) had to be uniformly distributed. The threshold was set to a minimum transition rate of 22% from one stimulus to the other. Passive listening trials were 2 s long (equal to the repetition time of the fMRI sequence). The auditory stimulus was presented between 1450 and 1550 ms after trial onset. The presentation of the auditory stimulus lasted 250ms. For additional details on the stimulus presentation see Preisig et al. (2021).

### 2.4 MRI data acquisition and preprocessing

Anatomical and functional MRI data were acquired with a 3-Tesla Siemens Prisma scanner using a 64-channel head coil. A 3-dimensional high-resolution T1-weighted anatomical volume was acquired using a 3D MPRAGE sequence with the following parameters: repetition time (TR) / inversion time (TI) / echo time (TE) = 2300/1100/3ms, 8° flip angle, FOV 256×216×176 and a 1×1×1 mm isotropic resolution. Parallel imaging (iPAT = GRAPPA) was used to accelerate the acquisition. The acquisition time (TA) of the T1-weighted images was 5 min and 21 sec.

Whole-brain fMRI data was acquired with sparse imaging to minimize the impact of EPI gradient noise during presentation of auditory stimuli (Hall et al., 1999). For this purpose, a delay was introduced in the TR during which the auditory stimuli were presented. This delay was 2070ms relative to the offset of the scan during task fMRI and 1070 ms during the passive listening run, respectively. For task fMRI each run included 128 echo planar imaging (EPI) volumes. The passive listening run included 336 EPI volumes. Each scan comprised 66 slices of 2mm thickness which were acquired using a interleaved acquisition sequence with multi-band acceleration (TR_task_: 3000 ms, TR_passive listening_: 2000 ms, TA: 930 ms, TE: 34 ms, flip angle: 90 deg, matrix size: 104×104×66, in-plane resolution: 2×2mm, Multi-band accel. factor: 6x). fMRI data were pre-processed in SPM12 (http://www.fil.ion.ucl.ac.uk/spm). For multivariate analyses, preprocessing included (I) functional realignment and unwarping and (II) co-registration of the structural image to the mean EPI. Univariate analyses included the additional preprocessing steps (III) normalization of the structural image to a standard template, (IV) application of the normalization parameters to all EPI volumes, and (V) spatial smoothing using a Gaussian kernel with a full-width at half maximum of 8 mm.

### 2.5 Multi voxel pattern analysis (MVPA)

The MVPA analysis was carried out in subjects’ native image space using realigned and unwarpped, but unsmoothed EPI images (Kriegeskorte et al., 2006). Voxel-wise BOLD activity was modeled by means of a single subject first-level GLM. The design matrix was specified including one regressor per condition and run coding for: unambiguous /da/ report, unambiguous /ga/ report, ambiguous /da/ report, and ambiguous /ga/ report. The regressors included all syllable reports for a given category, regardless of whether they were consistent or inconsistent with the presented stimulus. The onset of the button presses during task fMRI were modelled to account for the BOLD signal variability resulting from different response latencies. For each run, six realignment regressors accounting for movement-related effects and a constant term per functional imaging run were included in the model.

We conducted a whole-brain MVPA searchlight analysis (sphere radius 8mm, equivalent to 251 voxels) using the TDT toolbox (Hebart et al., 2015) to identify brain regions in which different syllable reports (/da/ vs /ga/) elicited distinct spatial BOLD response patterns. It is important to keep in mind that a perceptual identification task embeds perception in a context of decision- and response-making. Therefore, the crossnobis distance between /da/ and /ga/ reports may reflect not only the perceptual categorization process, but also decision making, motor planning, and the mapping of phoneme to finger. We further address this point in the discussion section.

For statistical inference, we computed the crossnobis distance between the response patterns associated with /da/ and /ga/ syllable reports within individual participants. The crossnobis distance is the cross-validated version of the Mahalanobis distance (multivariate noise normalized Euclidean distance) (Kriegeskorte et al., 2006; Walther et al., 2016), which results from an encoding approach (Hebart and Baker, 2018). To evaluate the representational consistency across unambiguous and ambiguous stimuli, the crossnobis distance between /da/ and /ga/ reports was computed as the arithmetic product of the distances within stimulus class (unambiguous and ambiguous stimulus trials).

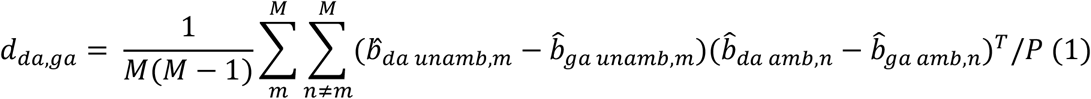

Where *d* denotes the crossnobis distance (in this case between /da/ and /ga/ syllable reports), *M* is the number of independent fMRI runs, 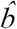 is the vector of responses to the condition consisting of the beta values of the searchlight voxels, and P is the number of voxels within the searchlight.

Like this, BOLD response patterns which consistently differentiated /da/ vs. /ga/ reports in both unambiguous and ambiguous stimuli yielded greater distances, while inconsistent BOLD patterns yielded smaller distances.

This analysis was implemented in the TDT toolbox (Hebart et al., 2015) using a encoding approach with leave-one-run-out cross-validation to estimate the crossnobis distance (Allefeld and Haynes, 2014). The cross-validation design was built using the *make_design_xclass_cv* function in the decoding toolbox (Hebart et al., 2015). The cross-validation design was specified such that the fitted regression coefficients (beta values) from the unambiguous trials were used for the training dataset and the beta values from ambiguous trials were used for the test dataset. In each of four cross-validation folds, the beta-value maps derived from unambiguous trials from three fMRI runs were cross-validated against the beta-value maps derived from ambiguous trials from the left-out run. Cross-validation ensures that the distance is zero if two voxel patterns are not statistically different from each other, making crossnobis distance a suitable summary statistic for group-level inference (e.g. with the one-sample t-test). Note that because of this cross-validation, the crossnobis distance can take negative values if its true value is close to zero in the presence of noise (Sohoglu et al., 2020).

For group-level inference, individual crossnobis distance maps were normalized and smoothed (using a Gaussian kernel with a full-width at half maximum of 8 mm) and then entered into a group level random-effects analysis using permutation-based nonparametric statistics in SNPM (http://www2.warwick.ac.uk/fac/sci/statistics/staff/academic-research/nichols/software). Unless stated otherwise, crossnobis distance maps from each participant were summarized at the group level using a non-parametric one-sample t-test against zero. False discovery rate (FDR) correction for multiple comparisons across voxels was applied at a threshold of p <.05.

### 2.6 Follow-up MVPAs

To further disentangle acoustic from phonetic representations, we tested whether categorical patterns derived from the unambiguous stimuli in the localized regions generalize better to the stimulus percept (/da/ vs /ga/) or to the presented acoustic stimulus (high vs low F3).

We would like to point out that this leads to a situation where the number of syllable reports that are inconsistent with the presented acoustic stimulus (e.g. /ga/ reports following high F3 and /da/ reports following low F3 stimuli) occur less frequently than the syllable reports that are in line with the F3 level (e.g. /ga/ reports following low F3 and /da/ reports following high F3 stimuli). What follows is an imbalanced number of trials per condition. Please note that crossnobis distance estimates the cross-validated multivariate noise-normalized means and therefore is not adversely affected by data imbalances (Hebart and Baker, 2018)

First, we recomputed the crossnobis distance between patterns associated with the syllable report within each stimulus class (high F3; low F3), hereafter referred (to as the MVPA Stimulus percept (/da/ vs. /ga/).

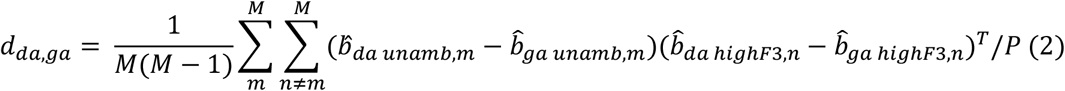

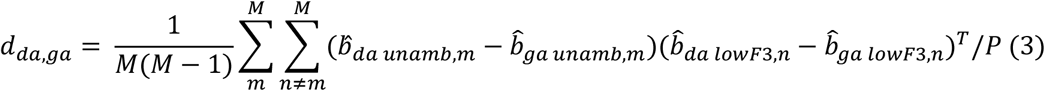

Thus, BOLD response patterns that differentiated /da/ vs. /ga/ reports consistently across unambiguous stimuli *and* within the same ambiguous stimulus yielded greater crossnobis distances, while inconsistent BOLD patterns yielded smaller distances.

The converse was also calculated – the distance between acoustic stimulus-evoked patterns, within each syllable report (/da/ vs /ga/), hereafter referred to as the MVPA Acoustic stimulus (high vs. low F3).

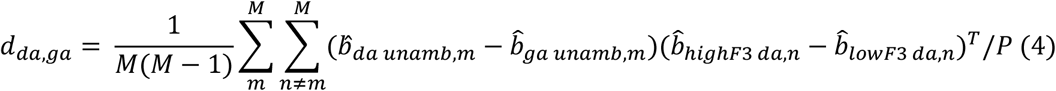

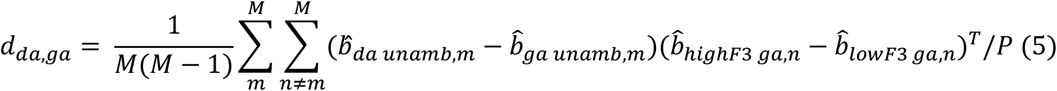

Thus, analogous to the above, BOLD response patterns that consistently differentiated unambiguous /da/ vs. /ga/ stimuli *and* high vs low F3 yielded greater distances, while inconsistent BOLD patterns yielded smaller distances.

For their data to be considered in one of the analyses (2-5), participants had to contribute at least two trials per fMRI run for each stimulus-response combination. The crossnobis-distance maps from participants who contributed to both analyses (2) and (3), or (4) and (5) were averaged. Data from 21 out of 27 participants was considered for each MVPA: Stimulus percept (/da/ vs. /ga/) and Acoustic stimulus (high vs. low F3). For group-level inference, we followed the same procedure that is described at the end of section 2.5.

### 2.7 Univariate analyses

In separate analyses, we mapped the brain activity from the passive listening fMRI run and the task fMRI runs. We decided to analyze the task and the passive listening condition in separate statistical models and qualitatively compare the descriptive results, because of different image acquisition procedures (different number of fMRI runs: one passive listening run vs. four task runs; different number of volumes: 336 per passive listening run vs 128 per task run; and different TR: 2s passive listening run vs 3s task run).

New first-level design matrices were specified using normalized and spatially smoothed images. Each design matrix (passive listening; task) included one regressor coding the onsets of all auditory stimuli per run. The task model contained an additional regressor coding the onset of the participants’ button presses. As before, each models included six realignment regressors to account for movement-related effects and a constant term per functional imaging run.

For each participant and model, T-contrasts (all auditory stimuli > implicit baseline) were computed to identify brain regions that responded significantly to auditory stimuli during passive listening and task (Pernet, 2014). Contrast maps from each subject and model were summarized at the group level using a one-sample t-test against zero.

We acknowledge that using separate statistical models limits our possibilities to draw statistical inference from the two conditions. Nonetheless, we believe that this qualitative approach allows us to describe which areas are active in the passive listening condition and which areas are active in the task condition.

## 3 Results

### 3.1 Binaural integration

Twenty-seven participants listened to ambiguous syllables (intermediate between /ga/ and /da/), whose perceived identity depends upon binaural integration, and to unambiguous syllables (clear /ga/ vs. /da/) whose categorization was unaffected by binaural integration (for details see Material & Methods) (Fig. 1). During task fMRI, participants reported on every trial whether they heard a /da/ or a /ga/ syllable.

Behavioral analyses indicated that participants reliably integrated the speech feature (high vs low F3) in the binaural integration condition. Participants answered on average with 25.40 ± 2.80% (mean ± SEM) /ga/ responses and 70.60% ± 3.00% /da/ responses to ambiguous syllables combined with the high F3 and 75.00% ± 3.5% /ga/ responses and 20.70% ± 3.10% /da/ responses for ambiguous syllables combined with the low F3. This is evidence for perceptual fusion of the dichotic stimulus, which is consistent with classical effects in dichotic syllable perception (Rand, 1974; Liberman and Mattingly, 1989; Preisig and Sjerps, 2019).

The probability of reporting an unambiguous stimulus with low or high F3 respectively as a /ga/ or /da/ syllable was high: Participants gave on average 84.4% ± 2.2% /da/ responses to unambiguous /da/ stimuli, and 87.4 ± 1.8% (mean ± SEM) /ga/ responses were registered for unambiguous /ga/ stimuli.

### 3.2 BOLD patterns differentiating /da/ vs /ga/ syllable reports

We conducted an MVPA searchlight analysis to identify BOLD response patterns which consistently differentiated /da/ vs. /ga/ reports in unambiguous and ambiguous stimuli. Group-level analysis revealed significant (*p* < .05 FDR-corrected) BOLD activity patterns in auditory and non-auditory brain regions that reliably represented the participants’ syllable report. One cluster extended from the anterior insula (AI) over the left inferior frontal gyrus (IFG), into the left ventral motor cortex (vMC). Another cluster was identified in the perisylvian region extending from the left supramarginal gyrus (SMG) into the left superior temporal gyrus (STG). A similar, but smaller cluster was also observed in the right hemisphere. Further, we found a cluster that included the left SMA and pre-SMA and a cluster that included part of the right somatosensory cortex (M1/S1) (see Fig. 2). The analysis also revealed significant clusters in the right anterior cingular cortex (ACC), the left superior frontal gyrus (SFG), the right cerebellum, the right middle frontal gyrus (MFG), and the left corpus callosum (CC) (see Table S1).

**Fig. 2.**
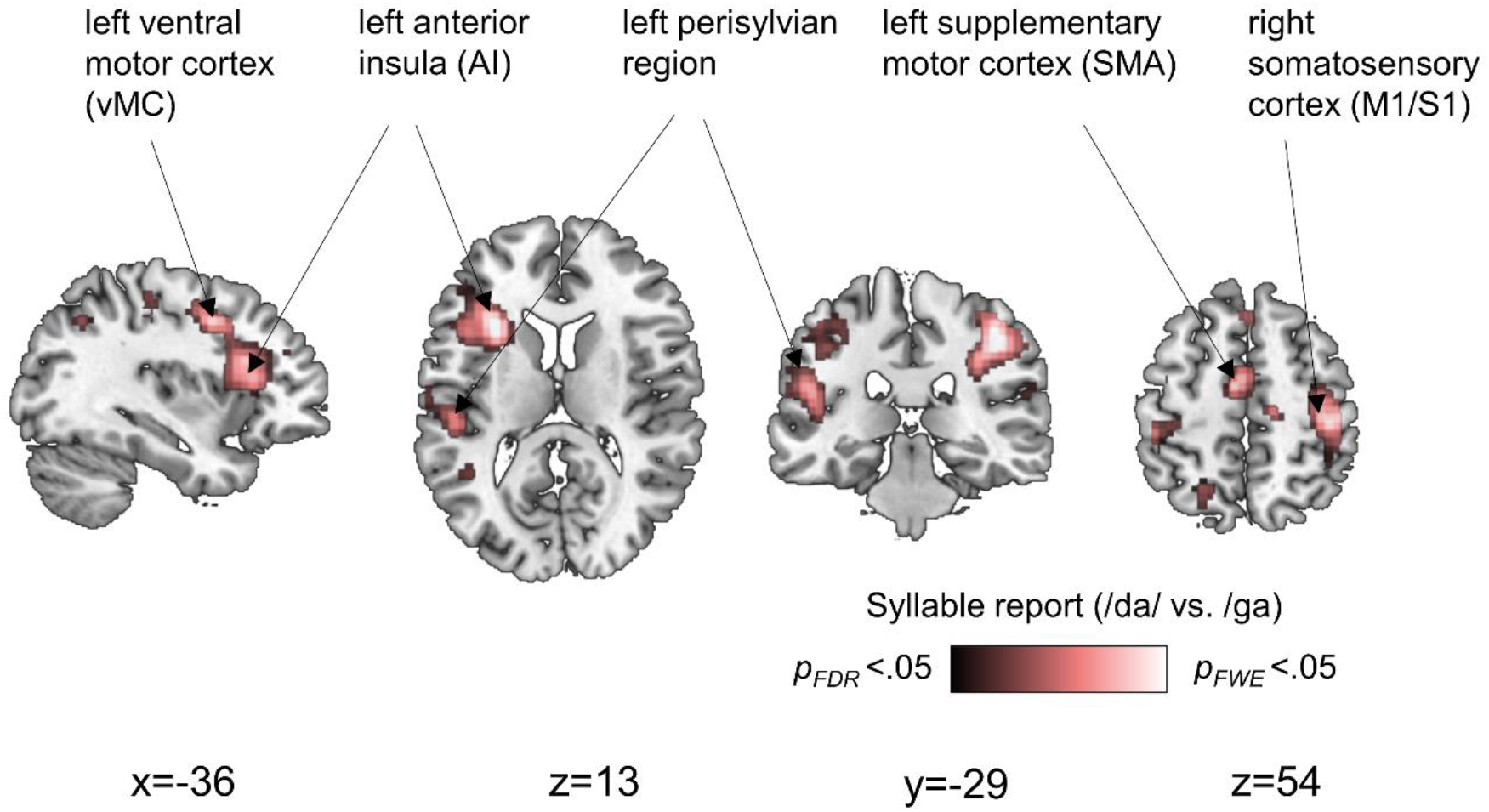
Results of the MVPA searchlight analyses projected onto a canonical MNI single-subject brain. In the highlighted regions, the average crossnobis distance associated with different syllable reports was found to be significantly larger than zero at *p*_*FDR*_ < .05. (B)

### 3.3 Acoustic or phonemic representation?

As physical stimulus and its perceptual interpretation were confounded in the MVPA analysis reported in section 3.2, the observed categorical response patterns could be driven by the stimulus acoustics, the syllable percept, or both. Thus, we tested in a follow-up analysis whether categorical patterns derived from the unambiguous stimuli in the localized regions generalize better to the stimulus percept (/da/ vs /ga/) within the same acoustic stimulus, or to the presented acoustic stimulus (high vs low F3) within the same stimulus percept.

For this purpose, we recomputed the crossnobis distance between syllable report-evoked patterns within each stimulus class (high F3; low F3). The converse was also calculated – the distance between acoustic stimulus-evoked patterns, within each syllable report (/da/; /ga/). In both cases the distances were cross-validated as previously, against the syllable reports in unambiguous stimuli.

A group-level analysis (random-effects analysis using permutation-based nonparametric statistics) of normalized and smoothed distance maps revealed significant BOLD activity patterns (*p* < .05 *FDR*-corrected) that differentiated between /da/ vs /ga/ percepts in the aforementioned areas (see Fig. 3A). In contrast, we found no significant clusters for the crossnobis distance maps between high vs low F3 acoustic stimulus at the same statistical threshold.

**Fig. 3.**
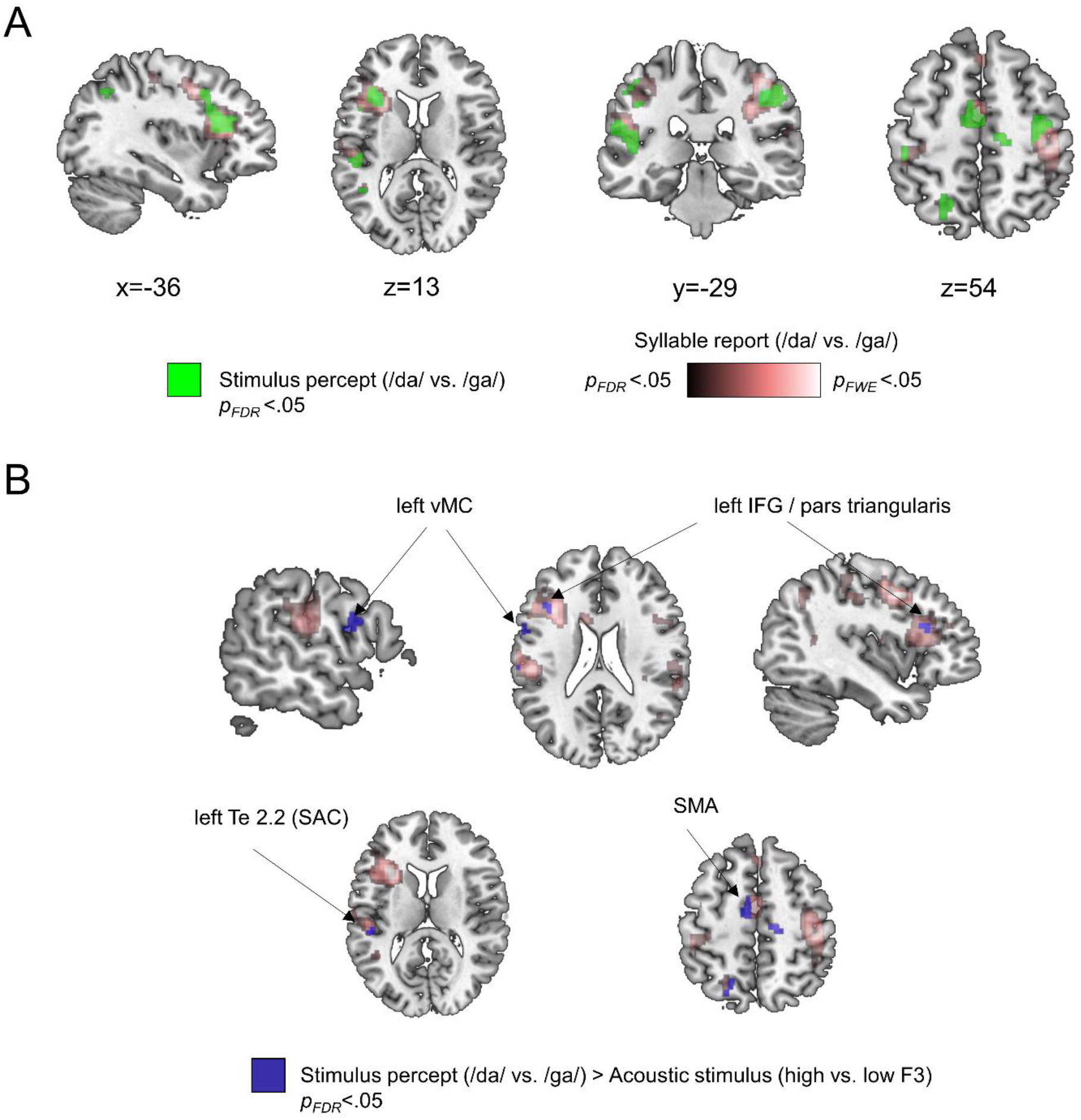
(A) Follow-up MVPA analysis constrained to the regions presented in Fig. 2. In green, crossnobis distance between /da/ vs /ga/ percept within the same acoustic stimulus (*p* < .05 *FDR*-corrected) is shown in green, overlaid on the clusters that discriminate different syllable reports across all trials. At the same threshold, we found no significant clusters for the crossnobis distance maps between high vs low F3 acoustic stimulus. (B) The contrast between perceptual and acoustic maps (paired t-test, *p* < .05 *FDR*-corrected) is shown in blue, overlaid on the clusters presented in Fig. 2.

In order to test for significant differences between syllable report-evoked patterns and acoustic stimulus-evoked patterns, we contrasted perceptual and acoustic maps at the group level (paired t-test using permutation-based nonparametric statistics). The analysis revealed that the crossnobis distance was significantly larger between different syllable percepts than between different stimulus acoustics in the left vMC, the left IFG (part triangularis), in the left Te 2.2, and the left SMA (see Fig 3B).

### 3.4 Auditory activation during passive listening and task

In order to see whether the identified syllable report-related regions co-localize with areas being activated by the syllable stimuli in the absence of any task and motor response, we mapped auditory-evoked activity during passive listening. For this purpose, we examined BOLD responses on all passive listening trials, contrasted with the baseline (see Methods).

During passive listening, whole-brain analysis revealed a bilateral brain network extending from the bilateral supratemporal plane into the IFG and the ventral AI. Further, we see activation in the right inferior parietal lobe extending into the right SMG and a cluster in-between the middle cingulum extending into the ventral pre-SMA (see Fig. 4, light blue color).

**Fig 4.**
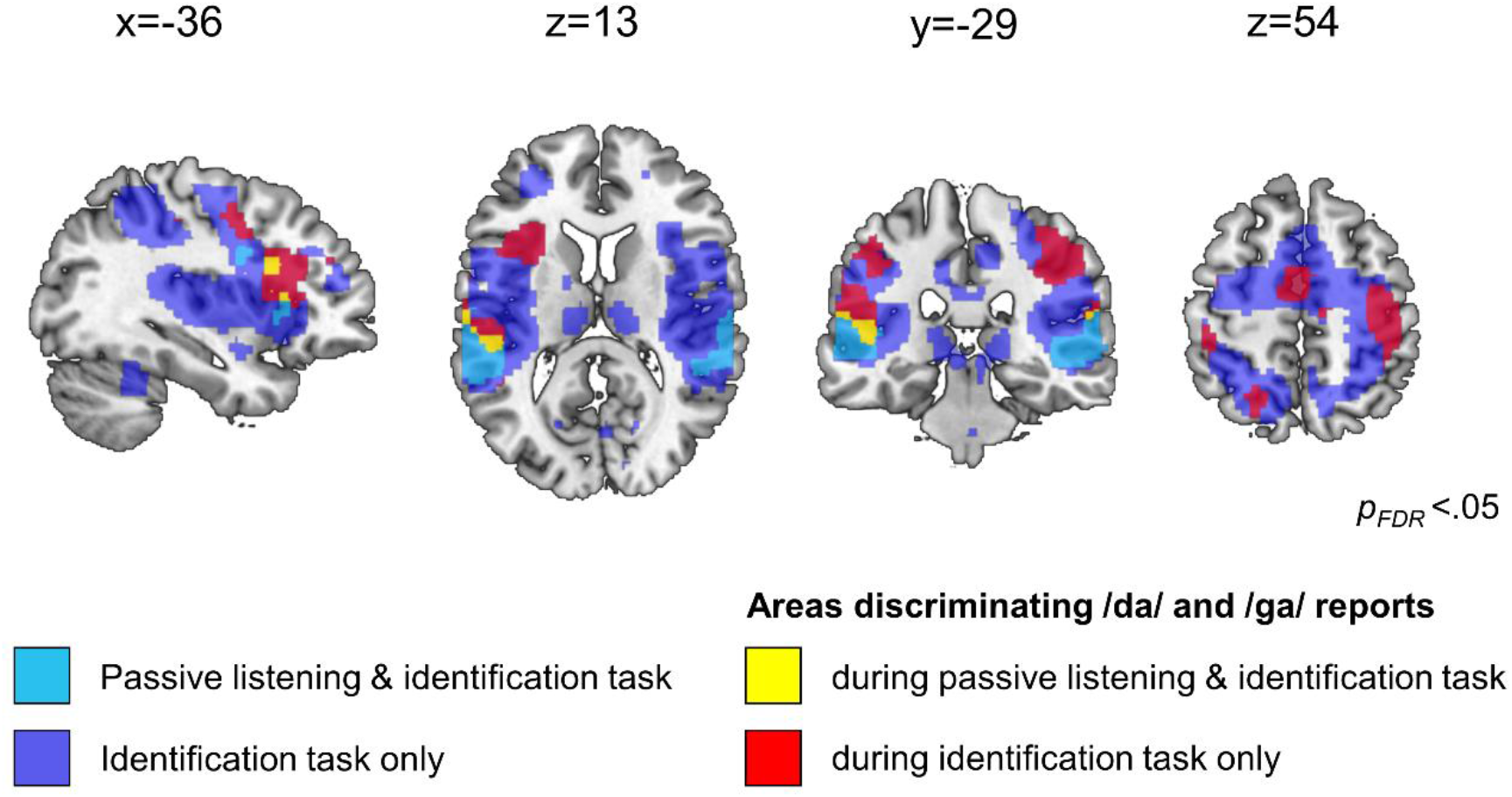
The clusters presented in Fig 2 overlaid on regions activated during auditory stimulus presentation and passive listening (light blue color) and regions activated during auditory stimulus presentation and task (dark blue color).

As illustrated in Fig. 4, regions that discriminate different syllable reports co-localize with areas activated during passive listening (see Fig 4, yellow color) in the left and right STG, in the left vMC, in the left inferior frontal operculum and the left AI. This suggests that processes unrelated to any task or motor response contribute to the neural representation that we identified.

Further, we also mapped the areas that were activated by the task to see whether syllable report-related regions co-localize with regions that are activated during the task, but not during passive listening (see Fig. 4). We examined BOLD responses on all task trials, contrasted with the baseline. In the task blocks, we observed more extensive activation in the same cortical areas and their surroundings as during passive listening (see Fig. 4, dark blue color). In addition, we observed activation in somatosensory, occipital as well as subcortical regions that was not present during passive listening.

Syllable report-related BOLD activity patterns co-localized in regions that were activated during the task, but not during passive listening, in the SMA and right M1/S1 (see Fig. 4, red color).

## 4 Discussion

In the present study, we aimed to identify the level of neural representation at which speech sounds are coded as abstract categorical perceptual units that are invariant to the sensory signals from which they are derived. For this purpose, we first localized brain regions which consistently discriminated the syllable reports evoked by different unambiguous and ambiguous stimuli. We then assessed whether these regions discriminated primarily the acoustics of the stimuli or the syllable report itself. Our results show that the syllable report was reflected in a set of auditory and non-auditory cortical regions which include regions in the bilateral perisylvian region (STG, SMG), left inferior frontal regions (vMC, IFG, AI), SMA/pre-SMA, and the right M1/S1. It is possible that the resulting lateralization of the neural activation reflects the lateralized presentation of the acoustic cue. However, binaural integration can occur independently of the ear to which the cue is presented (Preisig and Sjerps, 2019), thus it is unlikely that the lateralized presentation of the cue will affect the categorical neural response patterns. Future studies may confirm this by counterbalancing the ear of cue presentation. Importantly, most of these regions are outside of those traditionally regarded as auditory or phonological processing areas, such as the auditory cortex and superior temporal cortex. This result indicates that speech sound categorization implicates decision-making mechanisms and auditory-motor transformations. Further, it suggests the identified regions as a possible locus for transforming sensory signals into perceptual decisions.

When interpreting these findings, it is important to keep in mind that the task, as is typical of perceptual experiments, embeds perception in a context of decision- and response-making. We have divided the discussion of our results in four subsections below. Subsections 1-3 each represent a different hierarchy level: (1) auditory perception, (2) categorical perception of syllables, (3) motor response planning and execution. In subsection (4) we argue that there must be a level at which perception and motor planning must integrate to allow performing the task and that this integration involves also higher order cognitive processes such as decision making.

### 4.1 Auditory perception

One possibility is that the identified neural representation reflects auditory perception. Auditory speech perception is typically seen as a hierarchical process where simple acoustic features are extracted in early auditory areas (Schönwiesner and Zatorre, 2009; Leaver and Rauschecker, 2010; Norman-Haignere et al., 2015), which are later transformed into more complex and speech-specific representations in nonprimary cortex in the STG (Rauschecker and Scott, 2009; Brodbeck et al., 2018), possibly via parallel streams of auditory processing (Hackett et al., 2001; Jasmin et al., 2019; Hamilton et al., 2021). Indeed, the relatively subtle acoustic difference between our stimuli may suggest early auditory processing. However, we found no area where the crossnobis distance between different acoustic stimuli (high vs low F3) was larger than zero. It is possible that the acoustic difference was too subtle to be resolved from the fMRI data. Our analyses show that the identified BOLD patterns rather reflect the syllable percept than the acoustic difference between the stimuli. Therefore, we think it is likely that the identified BOLD patterns reflect processes beyond auditory stimulus perception.

### 4.2 Categorical perception of syllables

Another interpretation is that the identified patterns reflect the categorical perception of syllables. In line with existing data, we found that the bilateral pSTG/STS (Formisano et al., 2008; Chang et al., 2010; Kilian-Hütten et al., 2011; Mesgarani et al., 2014; Arsenault and Buchsbaum, 2015; Yi et al., 2019; Levy and Wilson, 2020), left SMG (Caplan et al., 1995; Dehaene-Lambertz et al., 2005; Zevin and McCandliss, 2005; Raizada and Poldrack, 2007) for a meta-analysis see (Turkeltaub and Branch Coslett, 2010), left IFG (Hasson et al., 2007; Myers et al., 2009; Lee et al., 2012; Chevillet et al., 2013; Du et al., 2014), the left vMC (Du et al., 2014; Evans and Davis, 2015; Cheung et al., 2016) and premotor cortex (Chevillet et al., 2013) discriminate between syllable reports in ambiguous and unambiguous stimuli. We found that all these areas also discriminate between different syllable reports when stimulus acoustics are kept constant. This effect was most visible in the left vMC, the left IFG (*pars triangularis*), and the left superior temporal lobe (Te 2.2), in which the crossnobis distances between syllable percepts were significantly greater than the distances between different stimulus acoustics (see Fig. 3B).

Further, the analysis revealed areas discriminating syllable reports that have rarely been associated with categorical speech perception: the left SMA and the left AI. In the midline motor areas, the strongest responses to auditory stimuli have been reported in pre-SMA and at the boundary area between pre-SMA and SMA (Lima et al., 2016). According to studies, which localized pre-SMA and SMA with fMRI (Mayka et al., 2006; Kim et al., 2010), the area identified in our study overlaps with the boundary area between pre-SMA and SMA. The SMA receives direct projections from the basal ganglia (Lehéricy et al., 2004; Akkal et al., 2007), the STG/STS (Luppino et al., 1993, 2001; Reznik et al., 2015), and inferior-parietal (Luppino et al., 1993) and inferior-frontal cortices (Catani et al., 2012; Vergani et al., 2014). The SMA is not typically considered as a part of the speech processing network in the brain (Hickok and Poeppel, 2007; Friederici, 2011; Hagoort, 2014), despite its connections with this network. Although SMA activity in response to speech and non-speech sounds has been reported in several studies (for reviews see Hertrich et al., 2016; Lima et al., 2016), the functional role of the SMA in auditory speech processing has remained elusive, possibly because SMA and pre-SMA are traditionally conceptualized as being linked to action-related processes, like speech motor control (Tourville and Guenther, 2011), rather than auditory processes (Lima et al., 2016).

The left AI cluster that we identified includes a portion of the dorsal anterior insula. Insula activity during speech perception has been reported in several studies of sublexical processing of speech (Benson et al., 2001; Golestani and Zatorre, 2004; Falkenberg et al., 2011) Particularly, the dorsal AI seems to be involved in speech perception, whereas other parts of the insula seem to contribute more to speech production (for a meta-analysis see Oh et al., 2014). Similar to the SMA, the insula has extensive connections with the auditory cortex, temporal pole, and superior temporal sulcus (Augustine, 1996; Oh et al., 2014), and is not typically considered in speech perception models (Hickok and Poeppel, 2007; Friederici, 2011; Hagoort, 2014).

Despite their contribution to auditory speech perception the SMA and the AI have rarely been associated with categorical speech perception. Possibly because previous studies either applied passive listening without task (Formisano et al., 2008; Lee et al., 2012), active listening with catch trials (Myers et al., 2009) or did not analyze brain signals from the SMA and the AI (Chang et al., 2010; Raizada et al., 2010; Kilian-Hütten et al., 2011; Mesgarani et al., 2014). Moreover, the contribution of motor action-related regions like the SMA may indicate that additional processes may contribute to speech sound categorization that go beyond categorical encoding as such.

### 4.3 Motor response planning and execution

Another explanation is that the identified response patterns reflect task-related motor or domain-general cognitive processes, such as button presses or response selection (Zatorre et al., 1992; Kawashima et al., 1996). This view is partially supported by our observation that the right M1/S1 differentiated between different syllable percepts, which we presume to be the result of the different motor patterns and tactile feedback for each response alternative. Nonetheless, it is likely that additional processes, other than responding, are responsible for the effect in these and other regions outside primary sensorimotor areas. Indeed, other regions in the identified network like the SMA, the left IFG or the left AI have been associated with motor planning of volitional movements (Tanji and Shima, 1994; Cunnington et al., 2005) and motor aspects of speech production (Ackermann and Riecker, 2004; Indefrey and Levelt, 2004; Oh et al., 2014). However, SMA (Benson et al., 2001; Scott et al., 2004; Warren et al., 2006; Jardri et al., 2007) and insula (Ackermann et al., 2001; Benson et al., 2001; Steinbrink et al., 2009; Hervais-Adelman et al., 2012) activation has also been reported during passive listening under various conditions. Further, there are studies showing involvement of SMA and insula during auditory tasks, which goes beyond merely motor-execution related activity, suggesting a role of these regions in accent imitation (Adank et al., 2013), vocal affect differentiation (Bestelmeyer et al., 2014), and categorization of different prosodic contours (statement vs question) (Sammler et al., 2015). Further, our results show that the regions which discriminate different syllable reports co-localize with areas activated during passive listening (in the left and right STG, in the left vMC, in the left inferior frontal operculum and the left AI). This suggests that other processes unrelated to any task or motor response contribute to the neural representation that we identified.

### 4.4 Speech sound categorization: A combination of different processes including higher order cognitive processes

The categorization of auditory information into syllables may involve various processes including auditory processing, categorical encoding and higher order cognitive processes such as decision making. In a perceptual task, decision making relies on sensory evidence (Gold and Shadlen, 2007). This decision process is not trivial because the sensory evidence provided by neural activation patterns is a mixture of signal and noise. Therefore, it would be an oversimplification to characterize speech sound categorization in terms of purely sequential stages of sensory decision making especially in higher-order cortical regions, given their strong top-down connections and involvement in recurrent processes.

A recent study found that SMA plays an important role for the transformation of auditory information into categorical responses during auditory decisions. Morán and colleagues (2021) recorded extracellular activity directly from SMA neurons in rhesus monkeys that were trained to categorize numerous complex sounds, including words. When the authors presented sounds along the continuum between two categories they observed that the neural response during auditory stimulus presentation varied gradually as a function of the changing sensory stimulus, while the neural responses during the button responses showed a categorical pattern that was consistent with the behavioral choices of the animal. This suggests that the SMA integrates acoustic information in order to form categorical signals that drive the behavioral response. Interestingly, the population activity in error trials reflected the behavioral decision, rather than the presented physical stimulus. In line with this findings, our follow-up analysis reported in section 3.3 revealed that the categorical patterns derived from unambiguous stimuli generalize better to the reported syllable (/da/ vs /ga/) than to the presented acoustic stimulus (high vs low F3). This indicates that higher order cortical regions influences may determine the activation patterns in auditory and somatosensory cortices.

It is possible that the left vMC has a similar function in speech sound categorization as the SMA. Skipper and colleagues (2007) investigated the role of the motor regions that are responsible for speech production during audiovisual speech perception. For their experiment, they used incongruent audiovisual stimuli (auditory /pa/ + visual /ka/) that elicit the phonemic percept /ta/ (Mcgurk & Macdonald, 1976). They found that “illusory” /ta/ percepts evoked activation patterns in the left vMC that were more similar to the activation pattern evoked by the audiovisual /ta/ stimulus than the activation pattern evoked by the audiovisual /pa/ or the audiovisual /ka/ stimulus. This suggests that left vMC activation represented the perceived stimulus rather than auditory or visual stimulus features. Most interestingly, the authors did an additional analysis where they compared the similarity of activations patterns over the entire time course of the hemodynamic response. For the left vMC, their analysis revealed that the activity patterns evoked by the “illusory” /ta/ percept showed the highest similarity with the activity evoked by the audiovisual /ta/ over the entire time course in the left vMC. However, the activation patterns in auditory and somatosensory cortices showed the highest similarity with audiovisual /pa/ for the first 1.5s of the hemodynamic response and were only afterwards more similar to the “illusory” /ta/ percept. This suggests that higher order cortical regions by means of top-down or recurrent processes may determine the activation patterns in auditory and somatosensory cortices and the phoneme percept as well as the behavioral response.

The left AI has been associated with decision making in a number of studies (for a review see (Droutman et al., 2015)). Particularly, the dorsal AI seems to be functionally implicated in cognitive control processes associated with decision-making, including attention re-focusing, evaluation, action, and outcome processing (Droutman et al., 2015). Further, dorsal AI activation has been related to decision ambiguity (Huettel et al., 2006) and error awareness (Ullsperger et al., 2010). Further recent evidence indicates that the anterior insula could serve as a gate for conscious access to sensory information (Huang et al., 2021).

We found two interesting distinctions between the BOLD patterns in the left SMA, left vMC, and the left AI. First, only the cluster in the left AI and left vMC overlapped with areas which were activated during passive listening to the same stimuli (Fig. 4). Second, only in the left SMA and the left vMC, but not in the left AI, the crossnobis distances between syllable percepts were larger than distances between different stimulus acoustics. This indicates that the responses in the left SMA and left vMC, but not left AI, were invariant to the physical stimulus properties. Taken together, one may speculate about the processing hierarchy and the functional differences between the processing in the AI, the vMC and the SMA. The AI may be more relevant for the integration of auditory evidence and the preparation of a perceptual representation, while the vMC and SMA serve to strengthen an acoustically invariant stimulus-syllable representation.

### 4.5 Conclusion

Our finding that BOLD patterns discriminating different syllable reports occur in brain regions contributing to auditory processing, categorical encoding and higher order cognitive processes suggests that the identified representations reflect a combination of various recurrent processes along this hierarchy.

Our finding that areas whose responses to speech stimuli discriminate between phonemic categories exist outside auditory cortical areas is consistent with the possibility that higher-order areas are instrumental in determining the syllables we hear, and that these regions feed abstract categorical representations back into the auditory association cortex (Formisano et al., 2008; Chang et al., 2010; Kilian-Hütten et al., 2011) and even to earlier auditory areas (Levy and Wilson, 2020).

## 5 Acknowledgement

This work was supported by the Swiss National Science Foundation [PZ00P1_201864 / PP00P1_163726] and the Janggen-Pöhn Stiftung. The authors would like to thank Benjamin Kop, Brigit Knudsen, Iris Schmits, Uriel Plones, and Paul Gaalman for their assistance as well as Martin Hebart for the methodological advice with regard to the MVPA analysis.

