## Supplemental material for "Speech sound categorization: The contribution of non-auditory and auditory cortical regions"

*Corresponding author: Basil C. Preisig, PhD

**This Document includes:**

Figures S1

Table S1

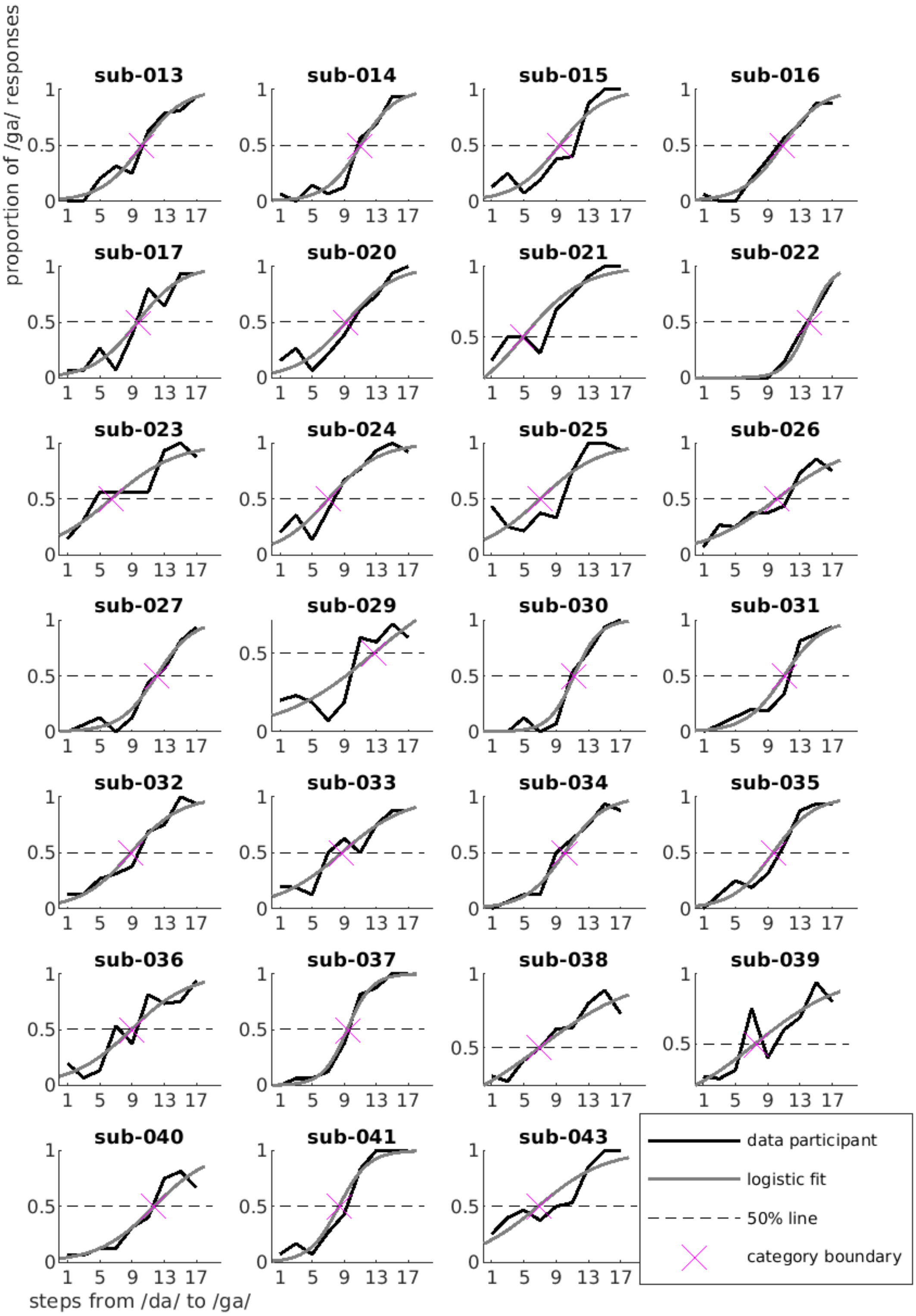


**Fig. S1. Psychometric curves of individual participants. The black solid line represents the proportion of /ga/ responses as a function of step from /da/ to /ga/. The grey line represents logistic fit. The horizontal dashed line is the 50% level. The magenta cross represents the estimated category boundary between /da/ and /ga/ reflecting the step at**

**which each participant reported perceiving the stimulus as /da/ or /ga/ in ~50% of the trials.**

**Table S1: The results of the group-level analysis (random-effects analysis using permutation-based nonparametric statistics) of normalized and smoothed individual distance maps (p< .05 $_{\text{FDR-corrected}}$)**

| | Coordinates | | | | |
|---|---|---|---|---|---|
| | x | y | z | T score | k |
| right postcentral gyrus (PoCG) | 42 | -26 | 44 | 6.47 | 2552 |
| left anterior insula (AI) | -30 | 20 | 12 | 6.25 | 2368 |
| left ventral motor cortex (vMC) | -38 | 2 | 42 | 5.51 | |
| left precentral gyrus (PCG) | -40 | 10 | 42 | 5.34 | |
| right anterior cingular cortex (ACC) | 4 | 34 | -4 | 5.49 | 141 |
| left superior frontal gyrus (SFG) | -22 | -2 | 66 | 5.48 | 75 |
| right cerebellum | 16 | -68 | -44 | 5.31 | 118 |
| left supplementarymotor cortex (SMA) | -6 | -8 | 54 | 5.05 | 817 |
| left pre-SMA | -6 | 12 | 62 | 3.26 | |
| left supramarginal gyrus / left superior temporal gyrus (STG) | -52 | -26 | 22 | 4.87 | 1588 |
| left postcentral gyrus (PoG) | -48 | -32 | 52 | 4.44 | |
| left angular gyrus | -40 | -60 | 42 | 3.72 | |
| superior medial frontal gyurs (at the border to the pre-SMA) | 0 | 28 | 50 | 4.68 | 244 |
| right superior medial frontal gyurs | 4 | 48 | 44 | 3.90 | |
| right middle frontal gyrus (MFG) | 22 | 44 | 32 | 4.62 | 198 |
| left corpus callosum (CC) | -10 | 14 | 24 | 4.16 | 107 |

Coordinates are in MNI space.
